# Structural Evolution of Neural Networks during Zebrafish Early Development

**DOI:** 10.64898/2026.10.29.762732

**Authors:** Zhengduo Wang, Li Tian, Bo Li

**Author notes:** **Correspondence and requests for materials** should be addressed to B.L. Corresponding author.

## Abstract

The spatial architecture of neural networks underpins nervous system function, yet dynamic *in vivo* evidence for their early establishment process remains scarce. Here, using long-term confocal live-cell imaging of transgenic zebrafish, we dissected neural network development across the brain, yolk sac, and spinal cord at single-cell resolution. We reveal that repeated growth cone bifurcation is the fundamental cellular mechanism for network node formation. Despite distinct developmental origins, networks in all three regions exhibit highly conserved dynamics, strictly following Strahler hierarchical ordering and Hack’s power-law scaling. Quantitative topological analysis shows that as development progresses, node numbers and clustering coefficients increase, while network density decreases. This reflects a structural optimization strategy balancing high functional efficiency with low wiring cost. These findings suggest that physical and geometric constraints, combined with simple local rules, drive the self-organization of complex neural networks, offering valuable principles for artificial network design.

## Introduction

The precise spatial architecture of neural networks constitutes the structural basis for all functional outputs of the nervous system (Luo, 2021). From peripheral circuits that regulate essential physiological activities such as heartbeat and reflexes, to the brain connectome that supports high-order cognitive functions including learning, memory and decision-making, the hierarchical organization, connectivity patterns and topological features of neural networks directly determine the efficiency, robustness and computational capacity of information processing (Bullmore & Sporns, 2012; Sporns, Tononi, & Kötter, 2005). Through hundreds of millions of years of evolution, biological neural networks have developed highly optimized construction principles: by adopting sparse connectivity, hierarchical branching and minimized wiring cost, they achieve exceptional functional performance under the constraints of limited spatial resources and energy supply (Bullmore & Sporns, 2012; Chklovskii, Schikorski, & Stevens, 2002; Hahn, Ponce-Alvarez, Deco, Aertsen, & Kumar, 2019). Such high-efficiency, low-cost structural design principles represent not only a core scientific issue in understanding neural development and evolution, but also provide important biologically inspired insights for the architectural optimization and energy efficiency enhancement of current artificial neural networks (Hassabis, Kumaran, Summerfield, & Botvinick, 2017; Jeon & Kim, 2023).

However, systematic *in vivo* dynamic evidence for the structural evolution of neural networks during early development remains scarce. This bottleneck arises because vertebrate embryonic development occurs *in utero*, and optically opaque brain tissue hampers continuous, real-time tracking of intact network formation at single-cell resolution (Lichtman, Livet, & Sanes, 2008; Nikolaou & Meyer, 2012). Traditional developmental neurobiology largely relies on fixed-tissue morphology, which provides only static snapshots at discrete time points and cannot resolve the quantitative laws governing continuous processes such as growth cone motility, branch formation, and topological evolution (Dent & Gertler, 2003; Rieger, Wang, & Sagasti, 2011). Although the zebrafish (*Danio rerio*) offers external embryonic development and whole-body transparency (Shen & Zuo, 2020), existing studies mostly focus on functional imaging of individual brain regions or specific neuronal populations (Migault, et al., 2018; Nelson & Granato, 2022). Consequently, a systematic investigation into the conserved rules governing neural network establishment across the brain, spinal cord, and yolk sac—as well as the quantitative principles underlying de novo network formation—remains lacking.

Building on our previous characterization of de novo neural development in zebrafish embryos (Wang, Tian, & Li, 2026b), we quantitatively dissected the topological features of neural networks here. By combining pan-neuronal fluorescent labeling with long-term confocal imaging, we tracked network establishment across the brain, yolk sac, and spinal cord at single-cell resolution. Using Tg (elavl3:EGFP) and Tg (elavl3:GCaMP6f) lines (Dunn, et al., 2016; Kim, et al., 1996; Park, et al., 2000), we documented the entire developmental process from initial axon outgrowth to complex interconnected networks. Through morphological node validation, Strahler analysis, and network topology quantification, we revealed that repeated growth cone bifurcation serves as the core mechanism for node formation. Despite distinct developmental origins and physiological functions, networks in all three regions exhibit highly conserved establishment dynamics and conform to Hack’s power-law scaling. As development progresses, these networks undergo topological optimization alongside increasing node numbers, displaying structural evolution marked by high functional efficiency and low wiring cost.

### Basic biological components and processes for network establishment

Using high-resolution confocal imaging, we observed that the establishment of the early neural network skeleton is essentially driven by the directed migration of growth cones. Axon extension is continuously pulled forward by the leading growth cone at its tip, and the dynamic motility of growth cones serves as the core driving force for both spatial expansion and branch formation of the entire neural network. As a dynamic structure at the axonal tip, the growth cone can undergo one or multiple rounds of bifurcation during its migratory extension. Following bifurcation of a single growth cone, two daughter axon branches are formed, and the bifurcation site thereby constitutes a node of the neural network, acting as a structural junction between adjacent axonal segments (**Fig. 1A; MovieS1**). At later stages of neural network development, growth cones originating from different neurons or distinct primary branches can also establish mutual contact and undergo anchoring fusion. This leads to cross-linking of previously independent axonal networks and the formation of compound nodes with higher node degrees, further promoting the increase in network connectivity complexity.

**Fig. 1.**
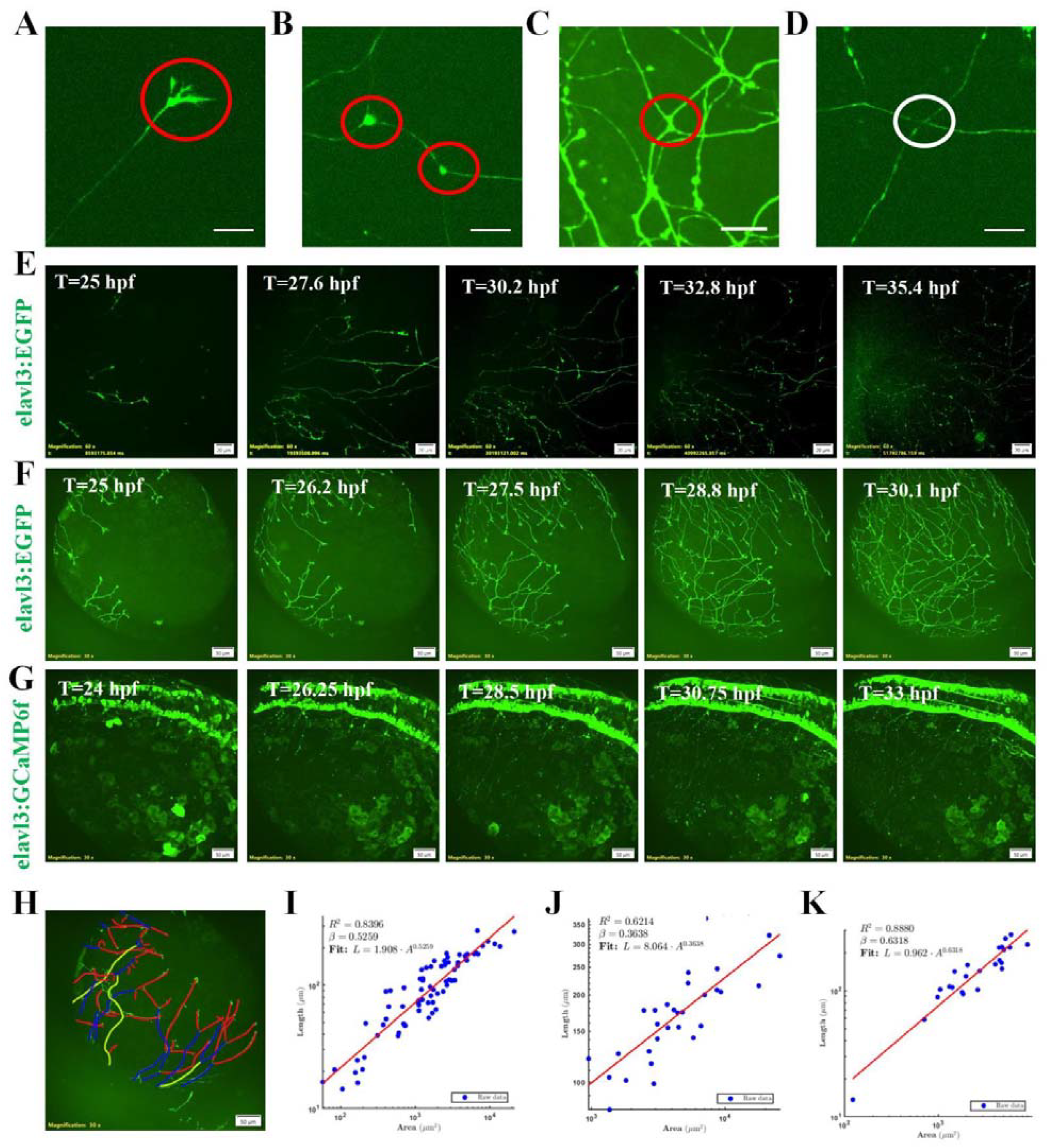
Dynamics of neural network formation in the brain, yolk sac, and spinal cord during zebrafish development. **(A)**, Representative confocal micrograph of neural network growth cones. Scale bar = 10 µm. **(B)**, Representative confocal micrograph of network nodes with a node degree of 3. Scale bar = 10 µm. **(C)**, Representative confocal micrograph of network nodes with a node degree of 4. Scale bar = 10 µm. **(D)**, Representative confocal micrograph showing axonal crossing without node formation in the neural network. Scale bar = 10 µm. **(E)**, Representative time-lapse confocal micrographs showing neural network growth in the brain of Tg (elavl3:EGFP) zebrafish embryos at 24 hours post-fertilization (hpf). Pan-neuronal signals are labelled with green fluorescence. Time-lapse imaging was acquired every 20 min using a 60× silicone oil immersion objective. Scale bar = 20 µm. **(F)**, Representative time-lapse confocal micrographs showing neural network growth on the yolk sac of Tg (elavl3:EGFP) zebrafish embryos at 24 hpf. Pan-neuronal signals are labelled with green fluorescence. Time-lapse imaging was acquired every 10 min using a 30 × silicone oil immersion objective. Scale bar = 50 µm. **(G)**, Representative time-lapse confocal micrographs showing neural network growth in the spinal cord of Tg (elavl3:GCaMP6f) zebrafish embryos at 24 hpf. Pan-neuronal signals are labelled with green fluorescence. Time-lapse imaging was acquired every 15 min using a 30× silicone oil immersion objective. Scale bar = 50 µm. Scale bar = 50 µm. **(H)**, Schematic diagram illustrating Strahler analysis for hierarchical architecture of the neural network. **(I)**, Power-law scaling between neural network covered area and maximum axon length in the zebrafish brain. **(J)**, Power-law scaling between neural network covered area and maximum axon length in the zebrafish yolk sac. **(K)**, Power-law scaling between neural network covered area and maximum axon length in the zebrafish spinal cord

When three-dimensional samples are rendered as maximum intensity projections (MIP), axons at different Z-planes may form intersections in the 2D projection plane, which can readily be misidentified as network nodes. We therefore established a discrimination criterion based on morphological features: authentic neural nodes (i.e., growth cone bifurcation sites) exhibit a characteristic triangular morphology, with one parent axon giving rise to two daughter branches. The fluorescence signal at the node is continuous and uniform, reflecting a true structural connection in three-dimensional space (**Fig. 1B, 1C; MovieS1**). In contrast, simple axonal crossings lack this triangular architecture; the fluorescence signals of the two overlapping axons represent merely spatial superimposition without structural continuity (**Fig. 1D; MovieS1**). This morphological criterion provides a reliable basis for node identification in subsequent quantitative network analyses.

We performed long-term time-lapse imaging on three anatomical regions—the brain, yolk sac, and spinal cord—to document the process of neural network establishment. In the brain, the neural network originates from clusters of neuronal somata. The somata extend axons tipped with growth cones outward; during extension, growth cones undergo repeated bifurcation, progressively forming a branched network. Axonal networks from distinct neuronal populations interweave with one another, ultimately developing into a complex cerebral neural network (**Fig. 1E; MovieS2**). In the yolk sac region, the neural network on the yolk sac surface is not generated by local neurons, but is derived from peripheral axons extending from neurons in the brain and spinal cord. These axons elongate and branch along the yolk sac surface, forming an expansive neural network covering the yolk sac (**Fig. 1F; MovieS2**). The developmental pattern of the spinal cord neural network is similar to that of the brain: axons extend from neuronal somata within the spinal cord and establish a branched network via growth cone bifurcation (**Fig. 1G; MovieS2**). The neural networks in all three regions follow the same fundamental developmental sequence: soma cluster, growth cone extension, bifurcation and branching and network formation, suggesting that this dynamic patterning mode is evolutionarily conserved during early neural development (Wang, Tian, & Li, 2026a).

### Conservative kinetic pathway across brain, spinal cord, and yolk sac

Given the striking morphological similarity between the branching patterns of neural networks and river drainage networks, we introduced the Strahler analysis method (Zhao, et al., 2025)—a geomorphic approach originally developed for stream ordering—to quantitatively dissect the hierarchical architecture of neural networks. The classification rules are defined as follows: the distalmost axons bearing growth cones are designated as order 1; when two axons of the same order converge at a node, the order of the resulting parent axon increases by one; when axons of different orders join, the resultant trunk axon retains the higher of the two orders (**Fig. 1H**). Strahler ordering clearly resolves the hierarchical organization of neural networks: lower-order axons correspond to peripheral growing branches responsible for territory coverage, whereas higher-order axons form the backbone pathways specialized for long-distance signal propagation. The neural networks in all three regions can be consistently hierarchically classified using this method, indicating that the topological logic underlying their branching architecture is evolutionarily conserved.

To explore the quantitative relationship between neural network extension length and coverage area, we defined network coverage area as the region enclosed by the growth cone fronts and the highest-order axons, and quantified the correlation between maximum axon length and the corresponding network coverage area in each region. The results revealed a significant power-law relationship between maximum axon length and network coverage area in all three anatomical regions—the brain, yolk sac, and spinal cord (**Fig. 1I–K**), consistent with Hack’s law in geomorphology (Dong, Vulis, Ma, Tejedor, & Goudge, 2026).

Specifically, the power-law exponents for the brain and spinal cord fell in the range of 0.5–0.6, indicating that neural networks in these two regions balance axial extension and areal expansion during development, such that network morphology maintains a relatively stable scaling as network size increases. The conserved presence of this power-law relationship across all three anatomical regions suggests that neural network growth may be governed by shared spatial constraints and optimization principles.

### Networking strategies over the course of development

Over the course of development, the number of neural nodes increases progressively in all three regions—the brain, yolk sac, and spinal cord—reflecting the continuous addition of branches and increasing structural complexity of the neural network (**Fig. 2A**). Quantification of node degrees (defined as the number of axons connected to a single node) revealed that the majority of nodes have a degree of 3 during development, corresponding to the three-branched structure formed by a single growth cone bifurcation. As development proceeds, the proportion of nodes with a degree of 4 increases gradually (**Fig. 2B, C**), indicating increased convergence and integration between distinct branch networks and a further elevation in the complexity of network connectivity. Analysis of the degree distribution showed that node degrees in the early neural network follow a skewed distribution: low-degree nodes predominate, while the number of high-degree nodes increases progressively with development (**Fig. 2D**).

**Fig. 2.**
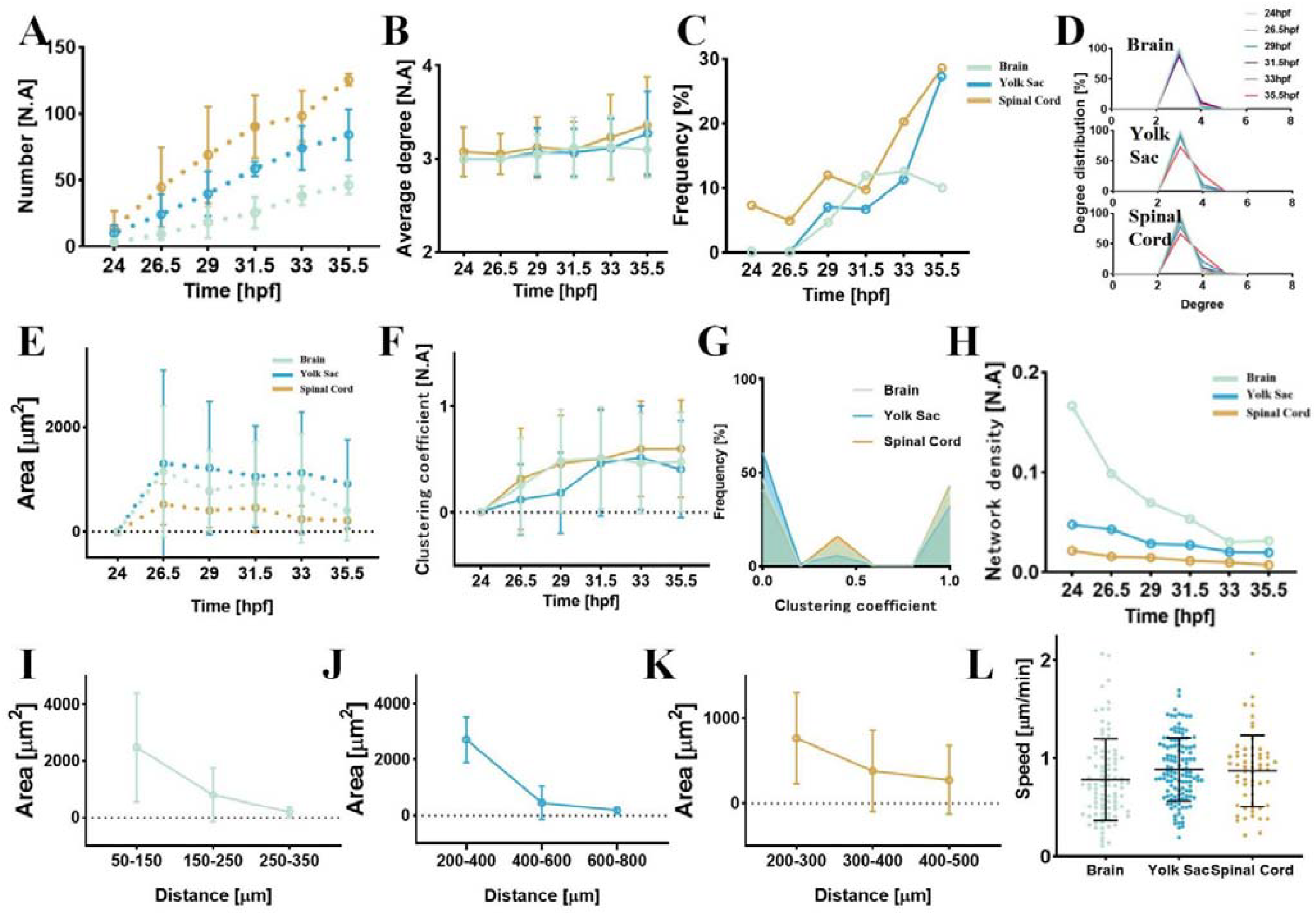
Quantitative analysis of neural network architecture in the brain, yolk sac and spinal cord during zebrafish development. **(A)**, Quantification of node number during neural network development in the brain, yolk sac and spinal cord. **(B)**, Line plot showing the average network degree of neural networks in the brain, yolk sac and spinal cord over developmental time. **(C)**, Frequency of nodes with a node degree of 4 in the developing neural networks of the brain, yolk sac and spinal cord. **(D)**, Degree distribution of neural networks in the brain, yolk sac and spinal cord at distinct developmental time points. **(E)**, Quantification of network void area during neural network development in the brain, yolk sac and spinal cord. **(F)**, Quantification of the clustering coefficient of neural networks in the brain, yolk sac and spinal cord over developmental time. **(G)**, Frequency distribution of clustering coefficients in neural networks of the brain, yolk sac and spinal cord. **(H)**, Quantification of network density of neural networks in the brain, yolk sac and spinal cord over developmental time. **(I)**, Quantification of distances between growth cones and voids of varying sizes in the developing brain neural network. **(J)**, Quantification of distances between growth cones and voids of varying sizes in the developing yolk sac neural network. **(K)**, Quantification of distances between growth cones and voids of varying sizes in the developing spinal cord neural network. **(L)**, Quantification of growth cone movement velocity in neural networks of the brain, yolk sac and spinal cord. For all statistical plots (n > 3 experiments) was conducted for all data. The error bars are the standard deviation of the data.

We used void area in MIP images as an indirect measure of the filling degree of the network. The results showed that void area in all three regions exhibits a decreasing trend with developmental progression, indicating a gradual increase in the coverage density of the neural network (**Fig. 2E**). Further analysis of the relationship between void size and distance to the growth cone front revealed that void areas are generally larger in the vicinity of growth cones, and decrease significantly in regions farther from the growth cones (**Fig. 2I–K**). This spatial distribution pattern indicates that refinement of the neural network is achieved through outward expansion of the leading growth cones and progressive space filling, and that the leading edge populated by growth cones represents the most structurally immature region of the network.

The clustering coefficient is a topological parameter that quantifies the tightness of local connectivity in a network. Statistical results showed that the clustering coefficient of neural networks in all three regions increases significantly over developmental time (**Fig. 2F**), indicating continuous improvement in the degree of local connectivity and integration. The frequency distribution of clustering coefficients exhibits a bimodal pattern, with the highest proportions at values near 0 and near 1, reflecting the concurrent presence of abundant peripheral branches (low clustering) and locally integrated nodes (high clustering) within the network (**Fig. 2G**). Notably, despite the continuous increase in node number, neural network density (the topological density of the network represented as an undirected graph, which measures the ratio of the number of actual edges to the theoretical maximum number of possible edges) exhibits a decreasing trend over the course of development (**Fig. 2H**). This result suggests that the early neural network does not undergo simple quantitative accumulation; instead, structural optimization occurs alongside branch addition (Wang, et al., 2026a, 2026b). By adjusting the spatial arrangement of branches, the network achieves more efficient coverage and connectivity at a lower wiring cost. This counterintuitive reduction in network density likely represents an evolutionarily tuned energy-saving developmental strategy. Early neural development requires widespread embryonic innervation, which relies on growth cone migration and bifurcation to achieve efficient spatial coverage. Concurrently, the network undergoes structural optimization during development to balance signaling efficiency and wiring cost, a trend most pronounced in the brain. In addition, we quantified growth cone migration velocity, which was comparable across the three regions and consistent with the network establishment rate in each respective region (**Fig. 2L**).

## Summary and discussion

By combining pan-neuronal fluorescent labeling with long-term *in vivo* confocal imaging at single-cell resolution, we systematically dissected neural network establishment across three distinct anatomical regions—the brain, yolk sac, and spinal cord—in early zebrafish embryos. We reveal that repeated growth cone bifurcation serves as the fundamental cellular mechanism driving network node formation and spatial expansion. Despite their distinct developmental origins and physiological functions, the networks in all three regions exhibit highly conserved developmental dynamics, strictly following Strahler hierarchical ordering and Hack’s power-law scaling. As development progresses, these networks undergo continuous topological optimization—characterized by increasing node numbers and local clustering coefficients alongside decreasing global network density—ultimately achieving high functional efficiency at low wiring costs.

These findings redefine the role of growth cone bifurcation, demonstrating that it acts not merely as a mode of individual axon branching, but as the foundational building block for the entire network architecture. The universal structural laws identified here strongly indicate that physical and geometric constraints, alongside genetic programs, significantly influence neural network establishment. Complex topologies can emerge from the coordinated effect of local rules—growth cone extension, bifurcating branching, spatial exclusion, and pruning—representing a highly efficient self-organizing mechanism that can substantially alleviate the burden of strict genetic encoding. Furthermore, the concurrent increase in local clustering and decrease in global density reveals an economical wiring strategy that emerges as early as the initial developmental stages, aligning with the wiring length minimization principle observed in mature brains. This provides biologically inspired insights for designing more efficient and lower-cost artificial neural networks with simpler algorithmic rules.

Several limitations remain to be addressed in future work. First, our MIP-based 2D projection analysis cannot fully capture the true three-dimensional spatial connectivity of the networks, necessitating further investigation incorporating advanced 3D reconstruction techniques and volumetric analysis. Second, this study focused exclusively on the structural establishment of axonal networks and did not cover synaptogenesis, which is critical for actual circuit function. Finally, the structure-function relationship of these networks awaits comprehensive validation via functional readouts, such as calcium imaging or electrophysiological recordings, to confirm how early structural evolution translates into functional maturation and information processing.

## Methods

### Construction and maintenance of transgenetic zebrafish lines

Zebrafish strains Tg(elavl3:GCaMP6f) was acquired from the China Zebrafish Resource Center (http://www.zfish.cn/), the Tg(elavl3:EGFP) was purchased from Nanjing Ezerinka Biotechnology Co., Ltd. (http://www.ezerinka.com/). All zebrafish were maintained in a recirculating aquaculture system procured from Nanjing Ezerinka Biotechnology Co., Ltd.

The system temperature was set to 28□°C, with continuous operation of UV sterilization and oxygen supply. A 5% sea salt solution and 5% sodium bicarbonate solution were used to regulate conductivity and ion balance, maintaining system conductivity between 500–800□μS/cm and pH between 7.0–8.0. Ambient temperature was controlled at 28□°C using air conditioning, and a 12□h light/12□h dark photoperiod was applied. Approximately one-third of the system water was replaced daily to keep total ammonia nitrogen below 0.02□mg/L.

To facilitate subsequent spawning, male and female zebrafish were housed separately. Males were distinguished by their slender body shape, flat abdomen, and lemon-yellow coloration, whereas females exhibited a fuller body, rounded abdomen, and silvery-gray pigmentation, allowing reliable visual sex identification. Adult zebrafish were fed live Artemia nauplii daily. The hatching procedure was as follows: 5□g of Artemia cysts and 5□g of sea salt were added to 1□L of pure water. The mixture was aerated under illumination for 24□h, after which aeration was stopped to allow eggshells to separate. The hatched nauplii settled at the bottom and were collected for feeding. Fish were fed 1-2 times per day, with the amount adjusted to ensure consumption within 15-20 minutes.

#### Sample preparation for *in vivo* zebrafish imaging

On the day before imaging, female and male zebrafish of different transgenic strains were placed in spawning tanks separated by a divider. The tanks were filled with a 1:1 mixture of fresh water and system water, covered, and maintained overnight. On the following morning, after the onset of illumination, the divider was removed to allow direct interaction between males and females. Males began nudging the abdominal region of the females, which subsequently initiated spawning. Approximately 30 minutes later, both females and males were returned to the main rearing system. The fertilized eggs were collected from the spawning tanks into 10 cm Petri dishes and maintained in specialized zebrafish embryo medium at 28□°C.

Embryo imaging typically commenced around 24 hours post-fertilization (hpf). Prior to imaging, embryos were screened using a spinning disk confocal microscope, and only those exhibiting clear fluorescence signals were selected for further processing. The chorion was then removed from each embryo to improve imaging quality. Although the chorion serves as a protective barrier during early development, it limits imaging depth and introduces motion artifacts as embryos become more active inside the membrane. For dechorionation, 24-hpf embryos were transferred to a Petri dish with a small amount of water and placed under a stereomicroscope. A small incision was made in the chorion using a corneal scissor, and the embryo was gently extruded with fine forceps.

The dechorionated embryos were then transferred to a confocal imaging dish. Low-melting-point agarose (0.12□g) was dissolved in 10□mL of ddH_2_O by microwave heating to prepare a 1.2% solution. After cooling to approximately 35□°C, the agarose solution was dispensed over the embryos. The position and orientation of each embryo were adjusted with forceps before the agarose solidified. For imaging at earlier developmental stages, such as 12 hours post-fertilization (hpf), the chorion was not removed due to the structural fragility of the embryos at this period, as mechanical manipulation could readily cause damage. Instead, the embryos within their intact chorions were directly embedded in low-melting-point agarose for immobilization during imaging.

#### Live cell imaging

Live imaging of zebrafish embryos was performed using a Spin SR10 spinning disk confocal microscope (Evident Olympus) equipped with an Oko-lab incubation chamber (H301-MINI). The system maintained a constant temperature of 28□°C throughout the experiments. Embryos were mounted in confocal dishes (Nest, 801001) and imaged with a typical laser intensity set at 20% and exposure time at 200□ms. For fluorescence excitation, the 488□nm laser line was used to visualize pan-neuronal labels in the transgenic lines (elavl3:EGFP, and elavl3:GCaMP6f).

To enhance experimental throughput, approximately 20 embryos were placed in each confocal dish. The microscope’s positional memory function was utilized to record individual embryo locations, enabling simultaneous time-lapse acquisition from multiple specimens. All embryos were imaged using axial Z-stack acquisition mode with the following parameters. Z-stack imaging was performed with a 60x oil objective at 100 μm Z-range and 2-3 μm step size, a 30x oil objective at 150 μm Z-range and 2-3 μm step size, and a 10x objective at 400 μm Z-range and 10 μm step size, respectively. Time-series imaging was performed at 20-minute intervals over a total duration of 48 hours, generating continuous 3D volumetric data throughout the recording period.

#### Data reconstruction

The reconstruction of imaging stacks from confocal microscopy is conducted using Cell Sens Dimension Desktop 4.2.1 (https://lifescience.evidentscientific.com.cn/en/software/cellsens/). In the top planar view, the Maximum Intensity Projection (MIP) algorithm is adopted to overlay all images in the z-stack into a single image. In the side view, the Marching Cubes (Surface Reconstruction Algorithms) algorithm is used to yield the stereoscopic spheroids. Compared with the range in the *x-y* plane, the length scale in *z* is relatively small, making the analysis based on *x-y* projection reasonably accurate.

#### Analysis of live cell imaging data

Time-lapse imaging data of zebrafish neural network development were analyzed using Fiji (ImageJ). Growth cone dynamics, including migration trajectories and velocity, were quantified with the Manual Tracking plugin. The analysis workflow consisted of the following steps: Time-lapse image stacks were loaded into Fiji, and the Manual Tracking plugin was initialized with appropriate temporal intervals and spatial calibration based on the acquisition settings. Growth cones were manually selected and tracked across consecutive frames using the plugin’s point-and-click interface. The plugin automatically recorded their positional coordinates over time and computed kinematic parameters such as displacement, velocity, and persistence.

The clustering coefficient is a topological metric that quantifies the extent of local connection clustering within a network, reflecting the tightness of connectivity in the neighborhood of individual nodes. For any given node *i* in the network, its clustering coefficient *C*_*i*_ is defined as the ratio of the number of actual edges present among all its adjacent nodes to the theoretical maximum number of possible edges between those neighbors, and is calculated as: 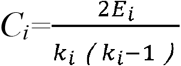,where *k*_*i*_ denotes the degree of node *i* (i.e., the number of edges directly connected to the node), and *E*_*i*_ represents the number of actual edges among the *k*_*i*_ adjacent neighbors of node *i*. The average clustering coefficient of the network is derived by calculating the arithmetic mean of the clustering coefficients of all nodes within the network. Meanwhile, the frequency distribution of clustering coefficients across all nodes is quantified to characterize the heterogeneity of local connectivity patterns in the network.

Graph density (i.e., topological density) is used to quantify the overall completeness of network connectivity, defined as the ratio of the number of actual edges to the theoretical maximum number of possible edges. The formula is given as: 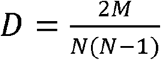, here *N* is the total number of nodes in the network, and *M* is the total number of actual edges present in the network. The term *(N(N-1)/2)* in the denominator corresponds to the theoretical maximum number of edges for an undirected graph with *N* nodes. Graph density was calculated independently for each of the three anatomical regions at different developmental time points, to characterize the dynamic changes in the global connectivity level of the network along the developmental progression.

## Acknowledgements

We thank Runjie Yu and Siyi Zhang from Oujiang Laboratory for assistance in spinning disk confocal and two photon microscopy imaging, respectively. DeepSeek V4 was used for text polishing under the supervision of the authors. This study was supported by taxpayers of China through National Natural Science Foundation of China (Distinguished Young Scholar Funding, Overseas; Young Scientists Fund-Category C, Grant No.32500692) and Wuhan University (Talents Startup Funding).

## Author contributions

B.L. conceptualized the project; B.L. Z.D.W. designed the research, Z.D.W, L.T. and B.L. performed experiments; Z.D.W. analyzed the data. All authors discussed the results and wrote the paper.

## Additional information

### Data availability

The data and analysis code that support the findings of this study are available upon reasonable request to B.L..

## Movie Captions

**Movie S1. Representative confocal time-lapse videos showing the assembly dynamics of the neural network in zebrafish embryos**. The first panel shows a representative confocal time-lapse video of growth cone migration pulling the expanding neural network. The second panel shows a representative confocal time-lapse video of growth cone splitting to generate two new axons and neural network nodes. The third panel shows a representative confocal time-lapse video of connection formation between neural nodes; neural network nodes are marked by red circles. The fourth panel shows a representative confocal time-lapse video of false nodes originating from maximum-intensity projection overlay of distinct axons; false neural network nodes are marked by red circles. A 30x silicone oil immersion objective lens was used. All neuronal structures were labeled with EGFP (excitation at 488 nm; laser power 20 %; exposure time 200 ms). Imaging started at 24 hours post-fertilization and was conducted every 10 minutes.

**Movie S2. Wide-field time-lapse confocal imaging was performed to record neural network development in the brain (left), yolk sac (middle), and spinal cord (right) of zebrafish (elavl3:EGFP; elavl3:GCaMP6f) embryos**. A 30x silicone oil immersion objective lens was used. All neuronal structures were labeled with EGFP (excitation at 488 nm; laser power 20%; exposure time 200 ms). Imaging started at 24 hours post-fertilization, with acquisition every 10-20 minutes for more than 14 hours. Scale bar = 20 or 50 μm.

## References

Bullmore, E., & Sporns, O. (2012). The economy of brain network organization. Nat Rev Neurosci, 13, 336–349.

Chklovskii, D. B., Schikorski, T., & Stevens, C. F. (2002). Wiring optimization in cortical circuits. Neuron, 34, 341–347.

Dent, E. W., & Gertler, F. B. (2003). Cytoskeletal dynamics and transport in growth cone motility and axon guidance. Neuron, 40, 209–227.

Dong, T. Y., Vulis, L., Ma, H., Tejedor, A., & Goudge, T. A. (2026). Apparent Hack’s law in river deltas. Science, 392, 493–498.

Dunn, T. W., Mu, Y., Narayan, S., Randlett, O., Naumann, E. A., Yang, C. T., Schier, A. F., Freeman, J., Engert, F., & Ahrens, M. B. (2016). Brain-wide mapping of neural activity controlling zebrafish exploratory locomotion. Elife, 5, e12741.

Hahn, G., Ponce-Alvarez, A., Deco, G., Aertsen, A., & Kumar, A. (2019). Portraits of communication in neuronal networks. Nat Rev Neurosci, 20, 117–127.

Hassabis, D., Kumaran, D., Summerfield, C., & Botvinick, M. (2017). Neuroscience-Inspired Artificial Intelligence. Neuron, 95, 245–258.

Jeon, I., & Kim, T. (2023). Distinctive properties of biological neural networks and recent advances in bottom-up approaches toward a better biologically plausible neural network. Front Comput Neurosci, 17, 1092185.

Kim, C. H., Ueshima, E., Muraoka, O., Tanaka, H., Yeo, S. Y., Huh, T. L., & Miki, N. (1996). Zebrafish elav/HuC homologue as a very early neuronal marker. Neurosci Lett, 216, 109–112.

Lichtman, J. W., Livet, J., & Sanes, J. R. (2008). A technicolour approach to the connectome. Nat Rev Neurosci, 9, 417–422.

Luo, L. (2021). Architectures of neuronal circuits. Science, 373, eabg7285.

Migault, G., van der Plas, T. L., Trentesaux, H., Panier, T., Candelier, R., Proville, R., Englitz, B., Debrégeas, G., & Bormuth, V. (2018). Whole-Brain Calcium Imaging during Physiological Vestibular Stimulation in Larval Zebrafish. Curr Biol, 28, 3723–3735.e3726.

Nelson, J. C., & Granato, M. (2022). Zebrafish behavior as a gateway to nervous system assembly and plasticity. Development, 149.

Nikolaou, N., & Meyer, M. P. (2012). Imaging circuit formation in zebrafish. Dev Neurobiol, 72, 346–357.

Park, H. C., Kim, C. H., Bae, Y. K., Yeo, S. Y., Kim, S. H., Hong, S. K., Shin, J., Yoo, K. W., Hibi, M., Hirano, T., Miki, N., Chitnis, A. B., & Huh, T. L. (2000). Analysis of upstream elements in the HuC promoter leads to the establishment of transgenic zebrafish with fluorescent neurons. Dev Biol, 227, 279–293.

Rieger, S., Wang, F., & Sagasti, A. (2011). Time-lapse imaging of neural development: zebrafish lead the way into the fourth dimension. Genesis, 49, 534–545.

Shen, C., & Zuo, Z. (2020). Zebrafish (Danio rerio) as an excellent vertebrate model for the development, reproductive, cardiovascular, and neural and ocular development toxicity study of hazardous chemicals. Environ Sci Pollut Res Int, 27, 43599–43614.

Sporns, O., Tononi, G., & Kötter, R. (2005). The human connectome: A structural description of the human brain. PLoS Comput Biol, 1, e42.

Wang, Z., Tian, L., & Li, B. (2026a). Neural Innervation Invigorates Yolk Sac Biological Functions beyond Nutrient Reservoir during Zebrafish Embryo Development. bioRxiv, 2026.2006.2006.730572.

Wang, Z., Tian, L., & Li, B. (2026b). Neurogenesis Leads Early Development in Zebrafish. bioRxiv, 2025.2011.2012.687769.

Zhao, A., Gruntman, E., Nern, A., Iyer, N., Rogers, E. M., Koskela, S., Siwanowicz, I., Dreher, M., Flynn, M. A., Laughland, C., Ludwig, H., Thomson, A., Moran, C., Gezahegn, B., Bock, D. D., & Reiser, M. B. (2025). Eye structure shapes neuron function in Drosophila motion vision. Nature, 646, 135–142.

